# Functional preference for object sounds but not for voices in the occipito-temporal cortex of early blind individuals

**DOI:** 10.1101/143776

**Authors:** Giulia Dormal, Maxime Pelland, Mohamed Rezk, Esther Yakobov, Franco Lepore, Olivier Collignon

## Abstract

Sounds activate occipital regions in early blind individuals. How different sound categories map onto specific regions of the occipital cortex remains however debated. We used fMRI to characterize brain responses of early blind and sighted individuals to familiar object sounds, human voices and their respective low-level control sounds. Sighted participants were additionally tested when viewing pictures of faces, objects and phase-scrambled control pictures. In both early blind and sighted, a double dissociation was evidenced in bilateral auditory cortices between responses to voices and object sounds: voices elicited categorical responses in bilateral superior temporal sulci while object sounds elicited categorical responses along the lateral fissure bilaterally, including the primary auditory cortex and planum temporale. Outside of the auditory regions, object sounds additionally elicited categorical responses in left lateral and ventral occipito-temporal regions in both groups. These regions also showed response preference for images of objects in the sighted, thus suggesting a functional specialization in these regions that is independent of sensory input and visual experience. Between-group comparisons revealed that only in the blind group, categorical responses to object sounds extended more posteriorly into the occipital cortex. Functional connectivity analyses evidenced a selective increase in the functional coupling between these reorganized regions and regions of the ventral occipito-temporal cortex in the early blind. In contrast, vocal sounds did not elicit preferential responses in the occipital cortex in either group. Nevertheless, enhanced voice-selective connectivity between the left temporal voice area and the right fusiform gyrus were found in the blind. Altogether, these findings suggest that separate auditory categories are not equipotent in driving selective auditory recruitment of occipito-temporal regions in the absence of developmental vision, highlighting domain-region constraints on the expression of crossmodal plasticity.

## Introduction

In visual and auditory areas of the human brain, separate brain clusters show category preferences. For instance, regions in the lateral part of the fusiform gyri (Fusiform Face Area or FFA) respond more to faces compared to other non-face objects (Kanwisher, McDermott, & Chun, 1997; Rossion, Hanseeuw, & Dricot, 2012). In contrast, non-face objects compared to faces typically elicit larger responses in parahippocampal gyri and in the latero-ventral aspect of the occipito-temporal cortex (Andrews & Schluppeck, 2004). Altough more seldom investigated than in vision, similar categorical preferences have been evidenced in the auditoy cortices so that listening to different categories of sounds such as humans voices (Belin, Zatorre, & Ahad, 2002; Belin, Zatorre, Lafaille, Ahad, & Pike, 2000; Kriegstein, Kleinschmidt, Sterzer, & Giraud, 2005) and artefacts (Lewis, Brefczynski, Phinney, Janik, & DeYoe, 2005; Lewis, Talkington, Puce, Engel, & Frum, 2011b; Lewis, Talkington, Tallaksen, & Frum, 2012) also activates distinct regions of the auditory temporal cortices. Importantly, a few studies have directly compared brain responses elicited by these sound categories while also taking into account their low-level characteristics, and suggest that categorical response in temporal cortex partially abstracts away from differences in basic sensory properties (Giordano, McAdams, Zatorre, Kriegeskorte, & Belin, 2013; Leaver and Rauschecker, 2010).

How sensory experience shapes these categorical preferences, particularly within the visual cortex, has recently received considerable attention. In people who lack visual experience because of early blindness, auditory and tactile stimulations massively activate the occipital cortex (e.g. Sadato et al., 1996; Weeks et al., 2000). Importantly, this reorganized occipital cortex is thought to follow a division of computational labour somehow similar to the one observed in the sighted (for reviews see Collignon, Voss, Lassonde, & Lepore, 2009; Dormal & Collignon, 2011; Reich, Maidenbaum, & Amedi, 2012; Ricciardi & Pietrini, 2011; Voss & Zatorre, 2012). For instance, dorsal occipito-parietal regions support spatial localization and motion processing in early blind subjects (Collignon et al., 2011; Dormal, Rezk, Yakobov, Lepore, & Collignon, 2016; Ricciardi et al., 2007; for reviews see Collignon et al., 2009; Dormal, Lepore, & Collignon, 2012), while occipito-temporal regions are responsive during tasks involving the identification of a non-visual input such as semantics and speech comprehension (Bedny, Pascual-Leone, Dodell-Feder, Fedorenko, & Saxe, 2011; Noppeney, Friston, & Price, 2003; Röder, Stock, Bien, Neville & Rösler, 2002), Braille reading (Büchel, Price, & Friston, 1998; Reich, Szwed, Cohen, & Amedi, 2011), or the discrimination of shape attributes of objects based on tactile (Amedi, Raz, Azulay, & Malach, 2010; Amedi et al., 2007; Pietrini et al., 2004), auditory (Amedi et al., 2007) or verbal material (He et al., 2013; Peelen et al., 2013). Results from these studies suggest that categorical organization of the VOTC may be observed, at least partially, independently of visual experience. Importantly, some authors have also observed similar domain preference in the VOTC of sighted individuals processing non-visual material, therefore suggesting that those regions may be a/meta/supra-modal by nature (Bi, Wang, & Caramazza, 2016; Heimler, Striem-Amit, & Amedi, 2015; Reich et al., 2012; Ricciardi, Handjaras, & Pietrini, 2014; Wang et al., 2015). In contrast, other studies have failed to show preferential responses to specific auditory categories in the VOTC of either blind (Mahon et al., 2009; He et al., 2013) or sighted people (Adam & Noppeney, 2010; Doehrmann, Naumer, Volz, Kaiser, & Altmann, 2008; Engel, Frum, Puce, Walker, & Lewis, 2009; Lewis et al., 2005; Tranel, Grabowski, Lyon, & Damasio, 2005). The questions our research aimed to address were straightforward: Do categories of sounds that show preferential activity in the temporal cortex specifically activate portion of the VOTC that typically represent similar categories visually ? Is visual experience influencing this functionally specific crossmodal activation of VOTC regions ? Are separate sound categories equipotent in inducing these functionally selective responses in VOTC ?

Categorical responses to sounds of objects *per se* that is, with non-verbal material and contrasting sounds of objects to another sound category, have only been investigated in sighted subjects so far. The few studies conducted in the sighted failed to identify a clear categorical selectivity in the VOTC (Adam & Noppeney, 2010; Doehrmann, Naumer, Volz, Kaiser, & Altmann, 2008; Engel, Frum, Puce, Walker, & Lewis, 2009; Lewis et al., 2005; Tranel, Grabowski, Lyon, & Damasio, 2005). Based on the numerous studies demonstrating that nonvisual (e.g. auditory) information elicits radically distinct patterns of responses in the occipital cortex of individuals with and without typical visual experience (Collignon et al., 2013; Laurienti et al., 2002; Sadato, Okada, Honda, & Yonekura, 2002) and that unique patterns of functional specialization (Bedny, Konkle, Pelphrey, Saxe, & Pascual-Leone, 2010; Collignon et al., 2011; Dormal et al., 2016; Weeks et al., 2000) and connectivity (Dormal et al., 2016) exist in early blind people, specific categorical responses to sounds of objects *per se* (i.e. not verbal material) could be expected in early blind individuals as a result of crossmodal plasticity (Bavelier & Neville, 2002).

In sighted humans, person (Mathias & Kriegstein, 2014), emotion (Collignon et al., 2008) and speech (van Wassenhove, 2013) recognition are all cognitive operations that benefit from the ability to efficiently bind facial and vocal stimuli (Yovel & Belin, 2013). It has recently been suggested that face-voice interactions might rely on direct functional and structural links between face selective regions in the visual cortex (e.g. Fusiform Face Area, FFA) and voice selective regions in the auditory cortex (e.g Temporal Voice Area, TVA) (Blank, Anwander, & Kriegstein, 2011). Preferential responses to voices in face selective regions, and *vice versa*, has however never been demonstrated in sighted individuals. In blind people, extracting crucial social information such as the speaker’s identity, emotional state and speech relies almost uniquely on voice perception. It could therefore be hypothesized that regions typically responsive to faces in the sighted would display a preferential response to voices in the blind, due to the enhancement/unmasking of pre-existing connections between these two cortical systems (Blank et al., 2011).

Here we used fMRI to characterize brain responses to object sounds, voices and their scrambled version in early blind and sighted individuals. We relied upon a factorial design that allowed us to directly contrast brain responses to familiar object sounds and to voices. The voice and object stimuli were controlled for global energy (Belin et al., 2000, 2002), but not for other low-level auditory cues that differ between voices and objects, such as the spectral content of the sounds (Belin et al., 2000, 2002). Control for these low-level cues is provided by the scrambling (see methods) of the two categories of stimuli. Therefore, by relying on directional contrasts between sound categories and between each category and its scrambled control condition, we assessed categorical preference while controlling for differences in the spectral content of the sounds and their global energy (see Rossion et al., 2012; Andrews, Clarke, Pell, & Hartley, 2010 for similar reasoning in the visual literature). Sighted individuals were additionally tested in a visual experiment involving pictures of faces, objects and scrambled pictures in order to assess the spatial correspondence between putative VOTC responses to auditory stimuli on the one hand, and categorical responses elicited by visual material on the other hand.

In summary, our goals were therefore threefold: (1) to test the existence of a double dissociation between voice and object selective-regions in temporal auditory cortices while controlling for differences in low-level properties of these sounds as in previous studies (Giordano, McAdams, Zatorre, Kriegeskorte, & Belin, 2013, Leaver and Rauschecker, 2010); (2) to investigate the existence of categorical responses to voices and sounds of objects in the VOTC and elucidate whether these putative responses are unique to the blind (due to crossmodal plasticity) or whether they are also observable in the sighted; (3) to explore differences in the functional connectivity profile of domain selective regions between blind and sighted people.

## Materials and Methods

### Participants

Thirty-three participants were recruited for this study. Sixteen early blind subjects (EB) (5 females, range 23 to 62 years, mean ± SD = 45 ± 12 years) (Table 1) and 15 sighted controls subjects (SC) matched to the EB group (5 females, range 22 to 61 years, mean ± SD = 41.9 ± 11.8 years) for age, gender, handedness, educational level and musical experience took part in the auditory experiment. Seventeen sighted participants, including the 15 that participated in the auditory experiment) were also tested in an independent visual experiment (7 females, range 22 to 61 years, mean ± SD = 40.7 ± 11.6 years). At the time of testing, the blind participants were totally blind or else had only rudimentary sensitivity for brightness differences and no pattern vision. In all cases, blindness was attributed to peripheral deficits with no neurological impairment (Table 1). All the procedures were approved by the research ethical and scientific boards of the “Centre for Interdisciplinary Research in Rehabilitation of Greater Montreal (CRIR)” and the “Quebec Bio-Imaging Network (QBIN)”. Experiments were undertaken with the consent of each participant.

**Table 1.** Characteristics of the blind participants. Handedness was evaluated using an adapted version of the Edinburgh inventory. Blind and sighted participants were classified as musicians if they had practiced a musical instrument or had vocal training for at least 2 years on a regular basis (at least 2 hours a week). A: Ambidextrous, M: male, F: female, m: months, y: years, OS: left eye, OD: right eye, Cegep: two years of education between High school and University.

| Participant | Age | Gender | Hand | Residual<br>Vision | Onset | Etiology | Educational<br>Level | Musical<br>Expe. |
| --- | --- | --- | --- | --- | --- | --- | --- | --- |
| EB01 | 48 | M | R | None | 1y | Glaucoma | University | Yes |
| EB02 | 44 | M | R | DL | 0 | Leber's congenital amaurosis | University | No |
| EB03 | 60 | F | R | None | 0 | Retinopathy of prematurity | High school | Yes |
| EB04 | 43 | M | R | None | 0 | Retinopathy of prematurity | High school | Yes |
| EB05 | 36 | F | R | None | 10m (OS)<br>/ 3.5y<br>(OD) | Retinoblastoma | Cegep | No |
| EB06 | 31 | M | R | None | 0 | Leber's congenital amaurosis | University | Yes |
| EB07 | 55 | M | R | None | 2m | Electrical burn of optic nerves | High School | No |
| EB08 | 51 | M | R | None | 0 | Glaucoma | University | Yes |
| EB09 | 45 | M | R | None | 0 | Retinopathy of prematurity | University | Yes |
| EB10 | 31 | F | A(R) | None | 0 | Retinopathy of prematurity | High School | No |
| EB11 | 51 | M | A(R) | None | 0 | Major eye infection | University | Yes |
| EB12 | 62 | M | R | DL | 0 | Congenital cataracts | Cegep | Yes |
| EB13 | 23 | M | R | DL | 0 | Glaucoma and microphthalmia | University | Yes |
| EB14 | 28 | M | R | None | 0 | Retinopathy of prematurity | University | Yes |
| EB15 | 57 | F | R | None | 0 | Chorioretinal atrophy (Toxoplasmosis) | Cegep | Yes |
| EB16 | 58 | F | R | None | 0 | Retinopathy of prematurity | Cegep | Yes |

### Experimental design and stimuli

Participants in both groups were scanned in an auditory run and were blindfolded throughout the fMRI acquisition. SC participants were additionally scanned in a visual run on a separate day. In order to familiarize the participants to the fMRI environment, participants underwent a training session in a mock scanner while listening to recorded scanner noise. Participants were familiarized to all stimuli before practicing the tasks in the mock scanner in order to ensure that all object sounds were clearly recognized by all participants. In the scanner, auditory stimuli were delivered by means of circumaural, fMRI-compatible headphones (Mr Confon, Magdeburg, Germany). Visual stimuli were projected on a screen at the back of the scanner and visualized through a mirror (127 mm × 102 mm) that was mounted at a distance of approximately 12 cm from the eyes of the participants.

#### Auditory stimuli

Auditory stimuli consisted of 4 different categories: human voices, object sounds, and their respective scrambled version (hereafter V, O, SV, SO, respectively) (Figure 1). All sounds were monophonic, 16-bit and sampled at 44.1 Hz. Voices and object sounds were cut at 995 ms (5 ms fade-in/fade-out). A 5 ms silence was added at the beginning of the stimuli to avoid them from clicking.

**Figure 1.**
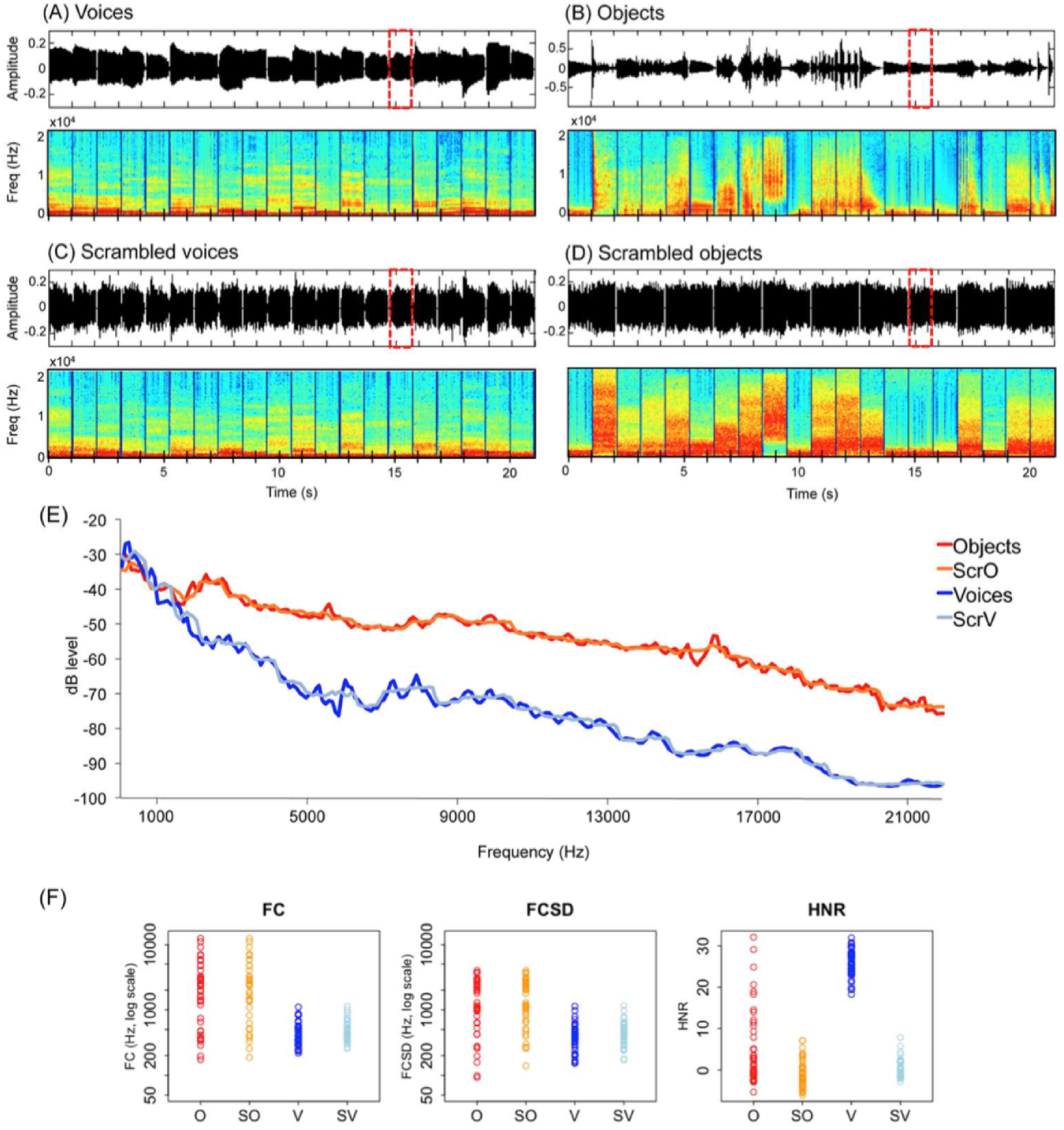
Stimuli in the auditory experiment. Top part: Sound properties in a representative 21 s block in the (A) voice and (B) object sounds conditions, and in the respective (C) and (D) scrambled conditions. Graphs represent sound amplitude as a function of time and frequencies spectrum as a function of time. Red dashed lines indicate the occurrence of a target sound (i.e repetition). (E) Bode magnitude plot expressing the magnitude in decibels as a function of frequency for the 4 blocks depicted in the top part of the figure. A sound block of each condition is available as supplemental material. (F) Measures of spectral content (FC and FCSD) and spectral structure (HNR) are plotted in color for each stimulus. Scrambling leaves the frequency spectrum relatively unaffected while altering harmonicity.

Human voices consisted of 8 exemplars of each of 5 vowels (“a”, “e”, “i”, “o”, “u”), pronounced by 40 different speakers (half male) recorded in the lab (Figure 1A). Object sounds consisted of 40 sounds of man-made artefacts (Figure 1B). In line with previous studies (Lewis et al., 2005; 2012; Lewis, Talkington, Puce, Engel, & Frum, 2011b), object sounds included a range of non-verbal sounds referring to non-living objects, namely human action sounds (lighting a match, jingling coins, hammering a nail, water flushing in the sink, jigsaw, manual saw, typing on a writing machine (2), dropping ice cubes in a glass, broom falling on the floor, pouring water in a glass, velcro, jingling keys, plate breaking, zipper, cleaning brush), bells and musical instruments (christmas bells, shop bell, door bell, piano, flute, drums, maracas, trumpet, guitar, tom tom, bycicle bell, harp) and automated machinery (car horn, train horn, helicopter, cuckoo clock, phone tone, motorcycle, gun bursts, printer, automatic camera, police car, tractor, hair dryer). These sounds were selected from a larger sample of 80 sounds in a pilot study based on the recognition performance of 10 sighted participants. In this pilot study, participants were asked to name each sound and subsequently rate it on a scale from 1 to 10 according to how much the sound was characteristic (representative) of the object. The 40 sounds with the highest rates (all above 7) were selected for the fMRI experiment. Before the actual fMRI experiment and prior to practicing the repetition task (see “Paradigm”) in the simulator, all participants were familiarized to each of the object sounds: they were asked to name each object after listening to its sound. Recognition accuracy during familiarization was at ceiling and was therefore not monitored.

Scrambled versions of the vocal and object sounds were obtained using MATLAB (The MathWorks, Inc., Natick, Massachusetts, United States) (Figure 1C and 1D). Scrambling was inspired by the method of Belin and colleagues (2000,2002) with the important difference that scrambling of amplitude and phase components was conducted separately within *frequency windows* (here 700 Hz) instead of *time* windows. Each vocal and object sound was submitted to a fast Fourier transformation and the resulting components were separated into frequency windows of ~700 Hz based on their center frequency. Scrambling was then performed by randomly intermixing the magnitude and phase of each Fourier component (Belin et al., 2000; 2002) within each of these frequency windows separately. The inverse Fourier transform was then applied on the resulting signal. The output was a sound of the same length of the original sound with similar energy within each frequency band (Figure 1A-D, power spectrum, and Figure 1E, dbLevel). For scrambled vocal sounds only, the envelope of the original voice was further applied on the output signal (Figure 1C). This was not done for scrambled object sounds because the application of the original envelope in this case lead many scrambled object sounds to be recognizable despite the scrambling (Figure 1D). Hence, for these sounds, a 5 ms ramp was applied in the beginning and at the end and a 5 ms silence was added at the beginning. Following standard practice, voices, object sounds and their scambled versions were equalized in root mean square (RMS) level (Belin et al., 2000; 2002; Giordano et al., 2013).

Measures of spectral content (FC and FCSD) and spectral structure (HNR) were extracted for each sound using Praat as described in Leaver and Rauschecker (2010) and are depicted in Figure 1F. FC reflects the center of gravity of the spectrum, an approximation of overall frequency content. FCSD is its standard deviation across the spectrum. HNR measures the ratio of the strength of the periodic and aperiodic (noisy) components of a signal.

The scrambling method used in the present study has the important advantage of altering the perception of the stimuli as object- and voice-like (see supplemental material for sound examples) while leaving the frequency spectrum of the original sound relatively unaffected (Figure 1). Temporal structure is relatively preserved only in the case of scrambled voices by application of the original sound envelope (Figure 1C). In contrast, harmonicity, typically higher for vocal stimuli, is altered by our scrambling method (Figure 1F, HNR).

This factorial design thus allows to control the frequency spectrum of objects and voices by contrasting these sounds to their scrambled versions. This is crucial considering recent evidence that occipital regions in congenitally blind subjects respond differently to distinct auditory frequencies (Watkins et al., 2013). Beyond controlling for low-level parameters of the sounds, this paradigm further allows to assess the degree to which low-level parameters contribute to a given categorical response (i.e. as it is the case for instance when a larger categorical response for voices relative to objects is also found when contrasting the corresponding scrambled control sounds, Table 2).

**Table 2.**
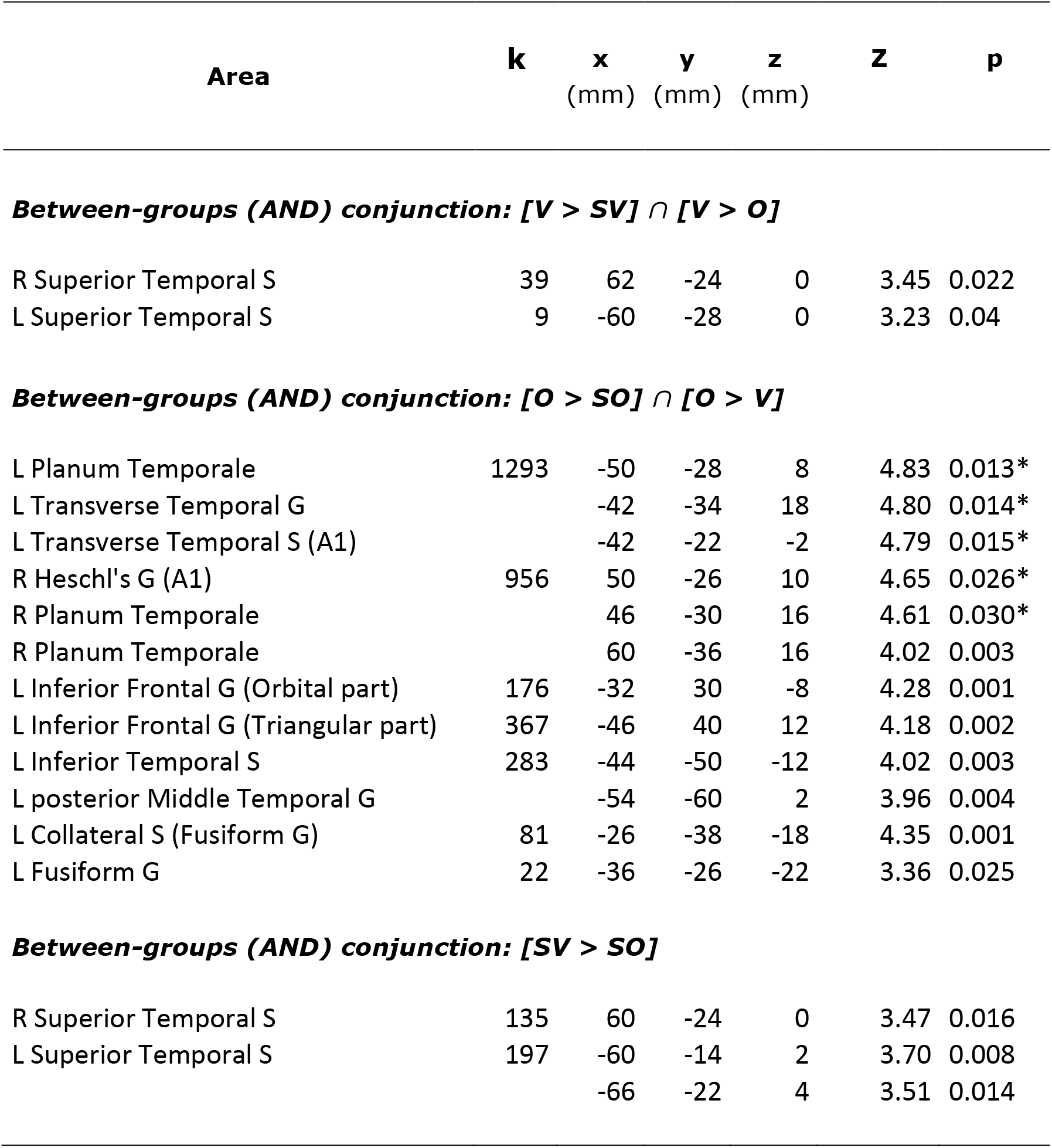
Categorical responses to voices and object sounds common to blind and sighted, and responses to low level properties of voices common to blind and sighted. Voice and object selective regions are depicted in Figure 2. Coordinates reported in this table are significant (p < 0.05 FWE) after correction over small spherical volumes (SVC) or over (*) the whole brain. K represents the number of voxels when displayed at p (unc) < 0.001. V: voices; O: objects; SV: scrambled voices; SO: scrambled objects; L = left; R=Right; G=Gyrus; S=Sulcus. Coordinates used for SVC are as follows (in MNI space): R Superior Temporal S: [60 −32 4] (Gougoux et al. 2009); L Superior Temporal S: [−64 −28 2] (Gougoux et al. 2009); R Planum Temporale: [52 −44 10] (Lewis et al. 2011); L Collateral S: [−28, −26 −26] (He et al. 2014); L Inferior Frontal G (Triangular): [−51 30 3] (Noppeney et al. 2003); L Inferior Frontal G (Orbital part): [−28 34 −6] (Bar et al. 2001); L Posterior MTG/ITG: [−52 −58 −6] (Peelen et al. 2013).

| Area | k | x<br>(mm) | y<br>(mm) | z<br>(mm) | Z | p |
| --- | --- | --- | --- | --- | --- | --- |
| **Between-groups (AND) conjunction: [V > SV] ∩ [V > O]** |  |  |  |  |  |  |
| R Superior Temporal S | 39 | 62 | -24 | 0 | 3.45 | 0.022 |
| L Superior Temporal S | 9 | -60 | -28 | 0 | 3.23 | 0.04 |
| **Between-groups (AND) conjunction: [O > SO] ∩ [O > V]** |  |  |  |  |  |  |
| L Planum Temporale | 1293 | -50 | -28 | 8 | 4.83 | 0.013* |
| L Transverse Temporal G |  | -42 | -34 | 18 | 4.80 | 0.014* |
| L Transverse Temporal S (A1) |  | -42 | -22 | -2 | 4.79 | 0.015* |
| R Heschl's G (A1) | 956 | 50 | -26 | 10 | 4.65 | 0.026* |
| R Planum Temporale |  | 46 | -30 | 16 | 4.61 | 0.030* |
| R Planum Temporale |  | 60 | -36 | 16 | 4.02 | 0.003 |
| L Inferior Frontal G (Orbital part) | 176 | -32 | 30 | -8 | 4.28 | 0.001 |
| L Inferior Frontal G (Triangular part) | 367 | -46 | 40 | 12 | 4.18 | 0.002 |
| L Inferior Temporal S | 283 | -44 | -50 | -12 | 4.02 | 0.003 |
| L posterior Middle Temporal G |  | -54 | -60 | 2 | 3.96 | 0.004 |
| L Collateral S (Fusiform G) | 81 | -26 | -38 | -18 | 4.35 | 0.001 |
| L Fusiform G | 22 | -36 | -26 | -22 | 3.36 | 0.025 |
| **Between-groups (AND) conjunction: [SV > SO]** |  |  |  |  |  |  |
| R Superior Temporal S | 135 | 60 | -24 | 0 | 3.47 | 0.016 |
| L Superior Temporal S | 197 | -60 | -14 | 2 | 3.70 | 0.008 |
|  |  | -66 | -22 | 4 | 3.51 | 0.014 |

#### Visual stimuli

In this experiment, stimuli consisted of 4 different categories: pictures of faces, objects and their phase-scrambled version (hereafter F, O, SF, SO, respectively) (Rossion et al., 2012). The *face* category consisted of full front pictures of 50 different faces (half male) (between 170 and 210 pixels width and 250 pixels height), that were cropped for external features and embedded in a white rectangle (220 pixels width × 270 pixels height). Similarly, the *objects* category consisted of pictures of 50 different objects (between 170 – 210 pixels width and 250 pixels height) inserted in a white rectangle (220 pixels width × 270 pixels height). The phase-scrambled pictures were used in order to control spatial frequencies and pixel intensity in each color channel (RGB) in the *face* and in the *object* categories. They were created using a Fourier phase randomization procedure by replacing the phase of each original image by the phase of a uniform noise allowing for amplitude to be conserved in each frequency band (Sadr & Sinha, 2004).

Pictures of objects consisted of the following items: fan, lamp, hat, garbage, coins, bag, balloon, stroller, glass, jeans, pair of boots, jewel, small bell, sofa, door, present, hairdryer, vase, hourglass, frame, headphones, key, clipboard, wine barrel, guitar, mug, toothbrush, tennis racket, alarm clock, tap, wardrobe, gloves, car tire, scissors, adjustable wrench, lens, screw, drum, trumpet, water gallon, light bulb, bucket, rugby ball, padlock, ring, paper bag, pepper, appel, plastic bag, ruby.

#### Paradigm

Both the auditory and the visual experiment consisted of a single run lasting about 18 minutes and consisting of 10 repetitions of each of the 4 conditions, alternating in blocks of 21 s. Blocks were separated by a 7 s baseline (silence and white fixation cross on a black background in the auditory and visual experiment, respectively). In each block, 20 items (sounds, pictures) were presented with a 50 ms inter-stimulus interval. Participants were instructed to detect a repetition in the stimuli (the same sound or picture presented twice in a row) by pressing a key with the right index finger. Emphasis was put on accuracy rather than speed. The number of repetitions within each block was unpredictable (i.e., 2 to 4 repetitions), thus ensuring that participants kept attending to the stimuli throughout the block. Within each condition, there were 4 blocks with one repetition, 4 blocks with 2 repetitions and 2 blocks with 3 repetitions, for a total of 18 targets/condition. This design aimed at matching as best as possible attention, arousal and motor components between conditions.

### Behavioral analysis

Behavioral performance in the auditory experiment was analyzed by submitting accuracy scores (hits minus false alarms) to a mixed 2 (*Group:* Blind, Sighted; between-subjects factor) × 4 (*Condition:* V, O, SV, SO) analysis of variance (ANOVA). In the visual experiment, a repeated-measures ANOVA was conducted with *Condition* (faces, objects, scrambled faces, scrambled objects) as a within-subjects factor. A Greenhouse-Geisser correction was applied to the degrees of freedom and significance levels whenever an assumption of sphericity was violated.

### MRI data acquisition

Functional MRI-series were acquired using a 3-T TRIO TIM system (Siemens, Erlangen, Germany), equipped with a 12-channel head coil. Multislice T2*-weighted fMRI images were obtained with a gradient echo-planar sequence using axial slice orientation (TR = 2200 ms, TE = 30 ms, FA = 90°, 35 transverse slices, 3.2 mm slice thickness, 0.8 mm inter-slice gap, FoV = 192×192 mm^2^, matrix size = 64×64×35, voxel size = 3×3×3.2 mm^3^). Slices were sequentially acquired along the z-axis in feet-to-head direction. The 4 initial scans were discarded to allow for steady state magnetization. The participants’ head was immobilized using foam pads. A structural T1-weigthed 3D MP-RAGE sequence (voxel size= 1x1x1.2 mm3; matrix size= 240x256; TR= 2300 ms, TE= 2.91 ms, TI= 900 ms, FoV= 256; 160 slices) was also acquired for all participants.

### Functional MRI analysis

Functional volumes from the auditory and the visual experiment were pre-processed and analysed separately using SPM8 (Welcome Department of Imaging Neuroscience, London, UK; http://www.fil.ion.ucl.ac.uk/spm/software/spm8/), implemented in MATLAB R2008a (The MathWorks, Inc., Natick, Massachusetts, United States).

Pre-processing included slice timing correction of the functional time series (Sladky et al., 2011), realignment of functional time series, co-registration of functional and anatomical data, creation of an anatomical template using DARTEL (a template including participants from both groups in the auditory experiment, and a template including sighted participants only in the visual experiment) (Ashburner, 2007), spatial normalization of anatomical and functional data to the template, and spatial smoothing (Gaussian kernel, 8 mm full-width at half-maximum, FWHM). The creation of a study-specific template using DARTEL was performed to reduce deformations errors that are more likely to arise when registering single subject images to an unusually shaped template (Ashburner, 2007). This is particularly relevant when comparing blind and sighted subjects given that blindness is associated with significant changes in the structure of the brain itself, particularly within the occipital cortex (J. Jiang et al., 2009; Pan et al., 2007; H.-J. Park et al., 2009).

#### Activation analyses

The analysis of fMRI data, based on a mixed effects model, was conducted in 2 serial steps, accounting respectively for fixed and random effects. In the audtitory experiment, changes in brain regional responses were estimated for each subject by a general linear model including the responses to each of the 4 conditions (V, O, SV, SO). These regressors consisted of boxcar function convolved with the canonical hemodynamic response function. The movement parameters derived from realignment of the functional volumes (translations in x, y and z directions and rotations around x, y and z axes) and a constant vector were also included as covariates of no interest. High-pass filtering was implemented in the design matrix using a cut-off period of 128 seconds to remove low-frequency noise and signal drift from the time series. Serial correlations in fMRI signal were estimated using an autoregressive (order 1) plus white noise model and a restricted maximum likelihood (ReML) algorithm. Linear contrasts tested the main effect of each condition ([V], [O], [SV], [SO]) and the contrasts between conditions ([V>O], [O>V], [V>SV], [O>SO]) and generated statistical parametric maps [SPM(T)]. These summary statistics images were then further spatially smoothed (Gaussian kernel 6mm FWHM) and entered in a second-level analysis, corresponding to a random effects model, accounting for inter-subject variance. For each of the abovementioned contrasts, one-sample t tests were performed within each group and two-sample tests were performed to compare effects between groups (EB>SC, SC>EB). Voice selective voxels were identified by means of a “AND” conjunction (Nichols et al., 2005) contrast of [V>O] and [V>SV]. Object selective voxels were identified by means of a conjunction contrast of [O>V] and [O>SO]. These contrasts thus identified voxels responding more to a category of sound relative to the other and for which this difference could not be accounted by differences in global energy or frequency spectrum.

These two conjunction analyses were conducted <u>separately for each group</u> (testing for voxels fulfilling these requirements in each group), jointly <u>between groups</u> (testing for voxels fulfilling these requirements in both groups, that is, independently of visual experience, see Figure 2A and 2B) and on between-group two-sample t-tests (testing for voxels fulfilling these requirements in one group more than in the other, Figure 3A).

**Figure 2.**
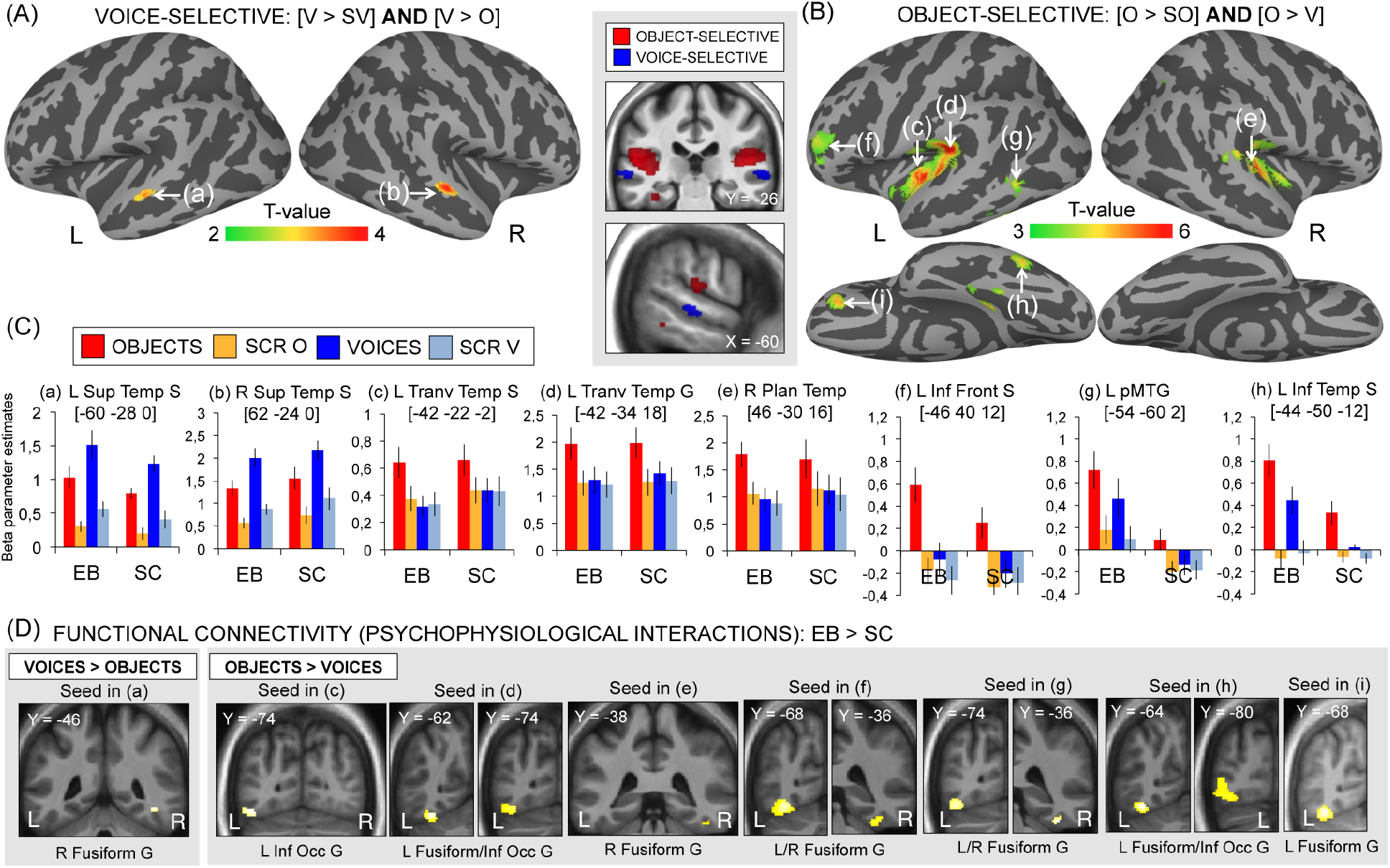
Categorical responses to (A) voices and (B) object sounds common to blind and sighted. For illustration purposes, activity maps are displayed at p(unc) < 0.005 with k > 90 (A) and p(unc) < 0.001 with k > 8 (B). Color bar represents t-values. (C) Mean activity estimates (arbitrary units ± SEM) are plotted for the 4 auditory conditions in significant peaks depicted in (A) and (B). (D) Psychophysiological interactions analyses as a function of group (blind > sighted) and experimental condition (V > O and O > V) based on the peaks of activation depicted in (A) and (B). For illustration purposes, activity maps are displayed at p(unc)< 0.005 and masked inclusively by the main effect in the blind (p(unc)< 0.005). EB = early blind; SC=sighted controls; L=left; R=right; S=sulcus; G=gyrus. See Table 2 and Table 6 for a list of brain regions depicted in this figure.

**Figure 3.**
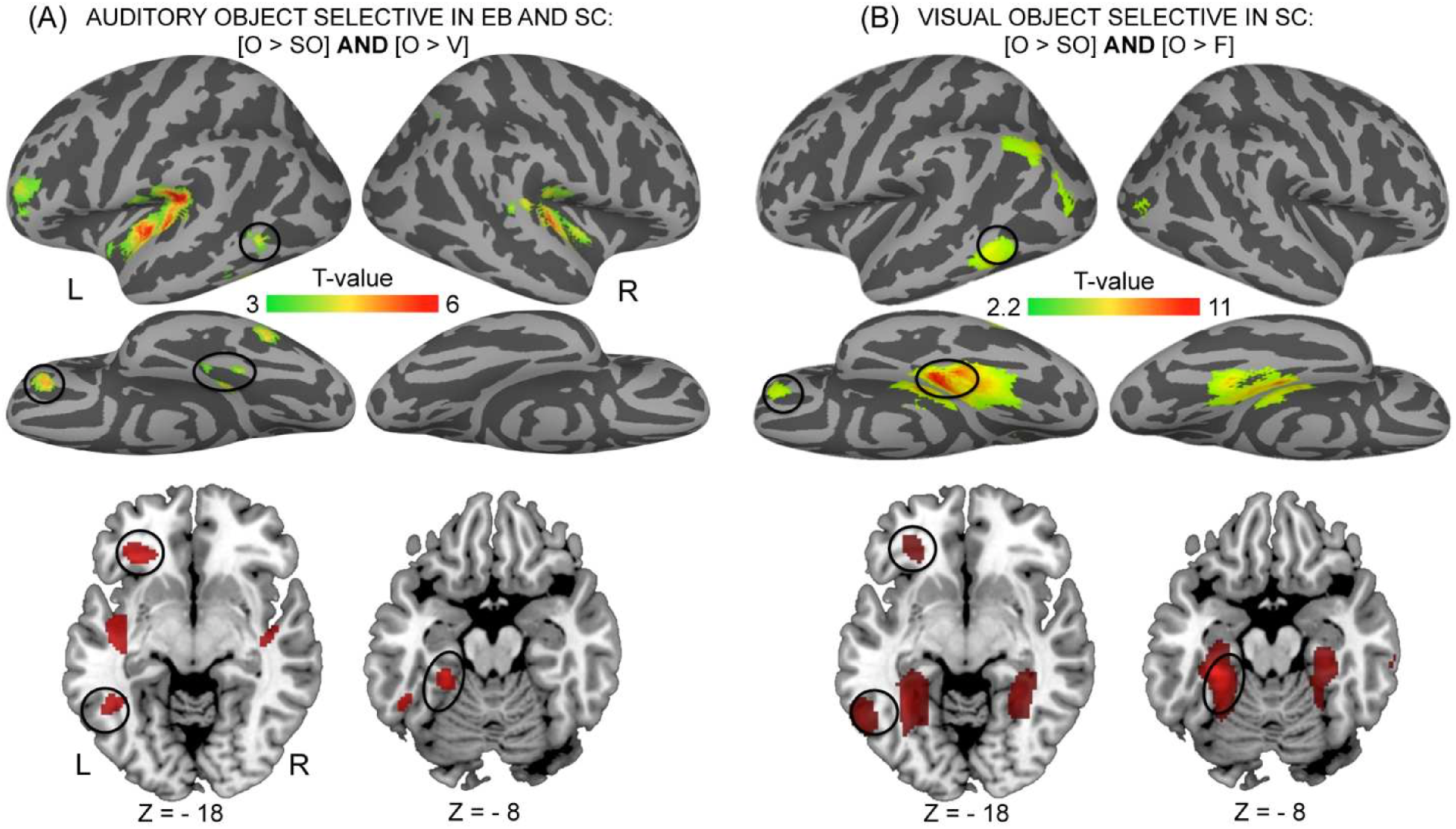
Objects categorical responses across modalities (visual and auditory) and populations (blind and sighted). (A) Categorical responses to object sounds common to blind and sighted. (B) Categorical responses to pictures of objects in the sighted. For illustration purposes, activity maps are displayed at p(unc) < 0.001 with k > 8. Color bar represents t-values. Regions in the left inferior frontal gyrus, posterior middle temporal gyrus and fusiform gyrus are responsive across groups in the auditory modality (A) and across the auditory and the visual modalities in the sighted (A and B). EB = early blind; SC=sighted controls; L=left; R=right; S=sulcus; G=gyrus; O = objects; V=voices; SO=scrambled objects; scrambled voices; F=faces; SF=scrambled faces. See Tables 2 and 3 for a list of brain regions depicted in this figure.

Preprocessing and statistical analysis of the fMRI data in the visual experiment were performed as in the auditory experiment with the exception that random effects were only calculated based on a one-sample t-test (no group comparison).

Statistical inferences were performed at a threshold of p < 0.05 after correction for multiple comparisons (Family Wise Error method) over either the entire brain volume, or over small spherical volumes (15 mm radius) located in structures of interest (see tables legends). Significant clusters were anatomically labeled using brain atlases (Petrides, 2012). Beta-weight extraction was used for visualization in figure charts only, while statistical analyses were performed on the single-voxel data, as per convention.

#### Psychophysiological analyses

Psychophysiological interaction (PPI) analyses were computed to identify any brain regions showing a significant change in functional connectivity with seed areas as a function of experimental condition (O, V) and group (EB>SC). Seed areas were selected using a two-step approach. First, all of the regions that were significant in the contrasts of interest, namely regions showing preferential responses to voices and objects in both groups (Figure 2) and those selectively responding to object sounds in the blind only (Figure 4) were selected as potential seed areas. Second, among these regions significantly active, seeds for PPI analyses were selected based on previous literature (see the list of selected regions in Table 6).

In each subject, the first eigenvariate was extracted using Single Value Decomposition of the time series across the voxels in a 10 mm radius sphere centered on the peak of activation reported at the group level. New linear models were generated using three regressors. The first 2 regressors were modeled as covariates of no interested and represented the condition (i.e. psychological regressor: O>V and V>O) and the raw activity extracted in the seed area (i.e. physiological regressor), respectively. The third, psycho-physiological regressor, represented the interaction of interest between the first (psychological) and the second (physiological) regressor. To build this third regressor, the underlying neuronal activity was first estimated by a parametric empirical Bayes formulation combined with the psychological factor and subsequently convolved with the hemodynamic response function (Gitelman, Penny, Ashburner, & Friston, 2003). Thus, variance explained by the psycho-physiological regressor is only that *above* variance explained solely by the main effects of task (psychological regressor) and physiological correlation (O’Reilly, Woolrich, Behrens, Smith, & Johansen-Berg, 2012). Movement parameters and a constant vector were also included as covariates of no interest. A significant PPI indicated a change in the regression coefficients between any reported brain area and the seed area, related to the experimental condition (O>V, V>O). Next, individual summary statistic images obtained at the first level (fixed-effects) analysis were spatially smoothed (6-mm FWHM Gaussian kernel) and entered in a second-level (random-effects) analysis using a one-sample t test. Two sample t-tests were then performed in order to compare these effects between groups.

Statistical inferences were performed as for the activation analyses with the exception that here we only report those regions showing a functional connectivity change in the blind compared to the sighted (EB >SC) and where the effect is driven by the blind. For this purpose, SVC (corrected for multiple comparisons using Family Wise Error method at p < 0.05) were performed on the between-groups functional connectivity maps (two sample t-tests thresholded at p (unc) < 0.001, EB>SC) and inclusively masked by the functional connectivity map in EB (one sample t-test, p(unc) < 0.001).

## Results

### Behavioral Results

#### Auditory experiment

There was no effect of group (p > 0.15), indicating that overall accuracy (hits – false alarms) did not differ between EB (mean ± SD = 91.58% ± 11.22%) and SC (mean ± SD = 86.2% ± 9.42%). There was a significant effect of condition (F_2.232,64.714_ = 4.493, p = 0.012) but group did not interact with this effect (p > 0.9). Two-tailed paired t-tests, collapsed across groups, revealed that detecting repetitions in the scrambled objects condition (mean ± SD = 83.87% ± 14.79%) was more challenging than in the other conditions (SV: mean ± SD = 90.86% ± 12.79%, t_30_ = −3.198, p = 0.003; O: mean ± SD = 92.29% ± 10.21%, t_30_ = −3.591, p = 0.001; V: mean ± SD = 88.89% ± 15.18%, t_30_ = −2.005, p = 0.05).

#### Visual experiment

There was a significant effect of condition (F_(3,48)_ = 9.663, p < 0.001). Two-tailed paired t-tests revealed that accuracy (hits minus false alarms) was lower in the scrambled faces condition (mean ± SD = 73.2% ± 21.26%) compared to the remaining conditions (faces: mean ± SD = 85.29% ± 11.44%, t_(16)_ = −2.766, p =0.014; objects: mean ± SD = 92.16% ± 9.43%, t_(16)_ = −4.788, p < 0.001; scrambled objects: mean ± SD = 87.58% ± 11.87%, t_(16)_ = −3.507, p = 0.003), and lower in the face than in the object condition (t_(16)_ = −2.159, p = 0.046).

### fMRI Results – Activation analyses

#### Object and voice-categorical responses common to early blind and sighted subjects

Between-group conjunction (AND) analyses identified brain regions commonly responsive in both groups when listening to voices compared to both scrambled voices and objects ([EB V > O] ∩ [EB V > SV] ∩ [SC V > O] ∩ [SC V > SV]), and when listening to objects compared to scrambled objects and voices ([EB O > V] ∩ [EB O > SO] ∩ [SC O > V] ∩ [SC O > SO]).

Categorical responses to voices common to EB and SC were found in 2 circumscribed areas within the superior temporal sulci bilaterally (Figure 2A, Table 2). Inspection of the individual data revealed that such responses were present in each single participant (data not shown). Importantly, these areas also strongly responded to low-level characteristics of voices (SV > SO, Table 2).

In both EB and SC, object sounds preferentially activated large portions of the auditory cortex bilaterally - although stronger in the left hemisphere - in the medial part of the transverse temporal gyrus (A1) extending laterally along the lateral fissure and posteriorly to the planum temporale. In the left hemisphere, additional clusters of activation were found within the inferior frontal gyrus and sulcus and within the temporal cortex, in the posterior middle temporal gyrus extending to the inferior temporal sulcus and fusiform gyrus (Figure 2B, Table 2). Contrary to voice-responsive regions, there was no contribution of low-level parameters to the response observed in object-responsive regions, neither in SC nor in EB (no significant responses in the contrast SO > SV). Object selective areas common to both groups in the left-lateralized inferior frontal gyrus, posterior middle temporal gyrus and fusiform gyrus overlapped with visual selective areas responsive to pictures of objects in SC (Figure 3B, Table 3) (as also confirmed by a conjunction analysis).

**Table 3.** Visually responsive regions in the sighted. Object selective regions are depicted in Figure 3B. Coordinates reported in this table are significant (p < 0.05 FWE) after correction over small spherical volumes (SVC) or over (*) the whole brain. K represents the number of voxels when displayed at p (unc)< 0.001. F: faces; O: objects; SF: scrambled faces; SO: scrambled objects; L = left; R=Right; G=Gyrus; S=Sulcus. Coordinates used for SVC are as follows (in MNI space): L Inferior Frontal G (Orbital part): [−28 34 −6] (Bar et al. 2001); L Posterior MTG/ITG: [−52 −58 −6] (Peelen et al. 2013), L Angular G: [−48 −70 31] (Fairhall and Caramazza, 2013).

| Area | k | x<br>(mm) | y<br>(mm) | z<br>(mm) | Z | p |
| --- | --- | --- | --- | --- | --- | --- |
| **Shape selective regions in vision: [O > SO]** |  |  |  |  |  |  |
| L Collateral S | 13013 | -34 | -34 | -18 | 5.76 | <0.001* |
| L Fusiform G |  | -46 | -54 | -18 | 5.41 | 0.002* |
| L Fusiform G |  | -38 | -46 | -10 | 4.96 | 0.015* |
| L Inferior Occipital G |  | -46 | -82 | -6 | 5.64 | 0.001* |
| L Inferior Occipital G |  | -56 | -66 | -14 | 4.98 | 0.013* |
| L Angular G |  | -48 | -70 | 24 | 4.77 | 0.030* |
| R Inferior Temporal G | 8510 | 50 | -70 | -10 | 5.53 | 0.001* |
| R Parahippocampal G |  | 32 | -24 | -22 | 5.52 | 0.001* |
| R Middle Temporal G |  | 52 | -74 | 4 | 5.24 | 0.005* |
| R Middle Temporal G |  | 58 | -14 | -18 | 5.30 | 0.004* |
| L Superior Frontal G | 5739 | -18 | -66 | 6 | 5.42 | 0.002* |
| L Inferior Frontal G (Orbital part) |  | -48 | 42 | -18 | 4.98 | 0.013* |
| L Inferior Frontal G (Orbital part) |  | -38 | 34 | -14 | 4.89 | 0.019* |
| L Inferior Frontal G (Orbital part) |  | -30 | 30 | -10 | 4.66 | 0.046* |
| L Superior Frontal G |  | -4 | 42 | 36 | 4.86 | 0.021* |
| **Object selective regions in vision: [O > SO] ∩ [F > SF]** |  |  |  |  |  |  |
| L Collateral S | 1295 | -32 | -34 | -18 | 5.75 | 0.000* |
| L Fusiform G |  | -30 | -50 | -14 | 4.67 | 0.044* |
| R Collateral S | 809 | 28 | -26 | -22 | 5.20 | 0.005* |
| R Fusiform G |  | 30 | -46 | -12 | 4.75 | 0.032* |
| L Angular G | 633 | -38 | -76 | 38 | 3.99 | 0.005 |
| L Angular G |  | -42 | -82 | 22 | 3.51 | 0.032 |
| L posterior Middle Temporal G | 360 | -58 | -58 | -6 | 4.29 | 0.002 |
| L Inferior Temporal G |  | -56 | -60 | -10 | 4.16 | 0.003 |
| L Inferior Frontal G (Orbital part) | 148 | -30 | 38 | -10 | 3.77 | 0.010 |

#### Object and voice-categorical responses specific to early blind subjects

Two-sample t-tests were then performed to compare these effects between groups. A conjunction (AND) analysis was conducted on the 2 sample t-tests [EB > SC] ×

[O > V] and [EB > SC] × [O > SO] in order to identify regions specifically activated in EB (relative to SC) for the processing of object sounds relative to both scrambled objects and voices (Figure 4, Table 4). This analysis revealed large bilateral activations in the occipital cortex that peaked in the middle and inferior occipital gyri bilaterally. There was no contribution of low-level parameters to the categorical response observed for objects (no significant responses in the contrast SO>SV, Figure 4B). These between-groups effects (EB>SC) were driven by the EB group (Table 4). The reverse group comparisons [SC > EB] did not reveal any region that was more strongly responsive in SC for object sounds relative to either voices or scrambled objects.

**Figure 4.**
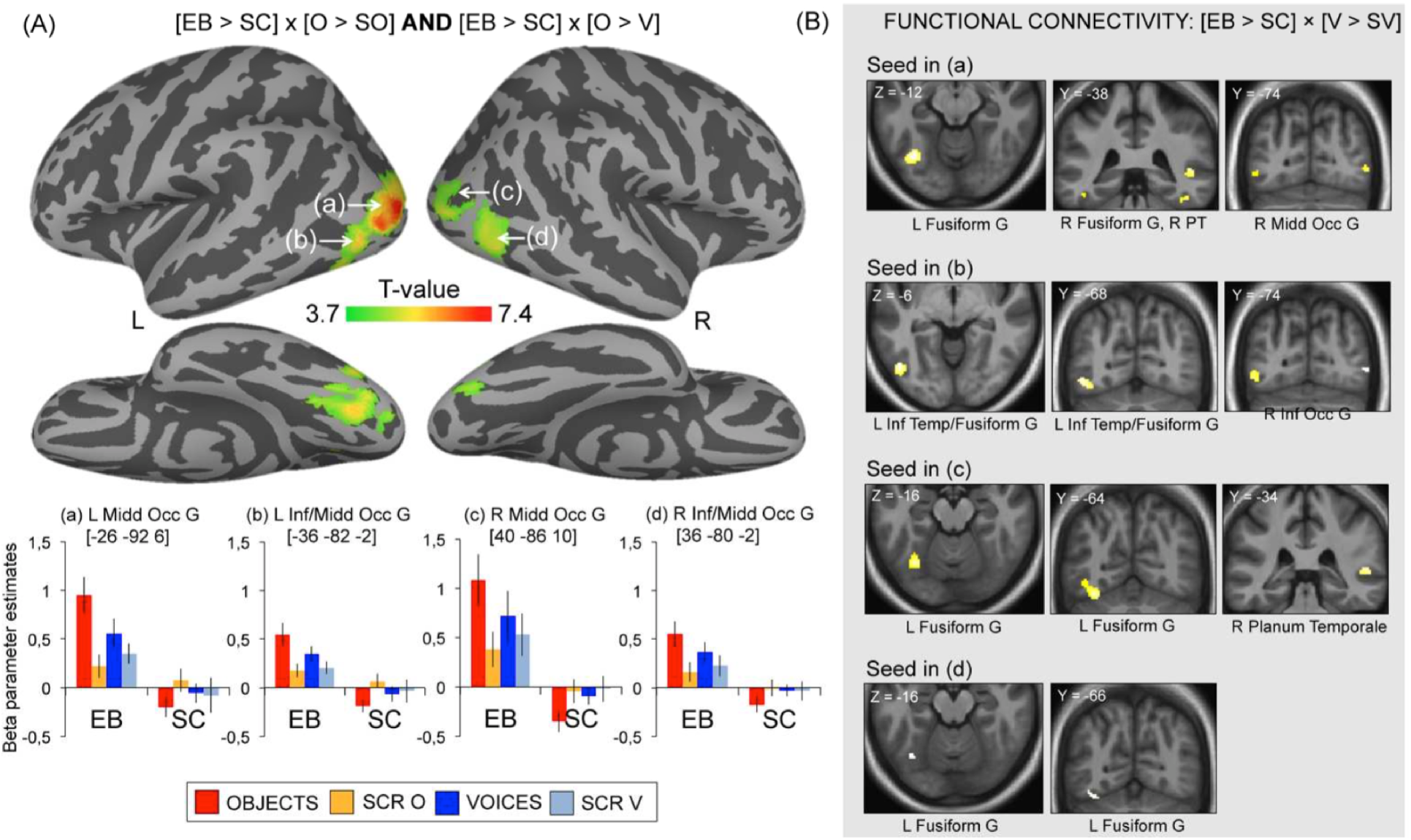
Categorical responses to object sounds specific to the blind. Color bar represents t-values. For illustration purposes, activity maps are displayed at p(unc) < 0.0001. (B) Mean activity estimates (arbitrary units ± SEM) are plotted for the 4 auditory conditions in significant peaks depicted in (A). (C) Psychophysiological interactions analyses as a function of group (blind > sighted) and experimental condition (O > V) based on the peaks of activation depicted in (A). For illustration purposes, activity maps are displayed at p(unc) < 0.001 and masked inclusively by the main effect in blind group (p(unc) < 0.001). See Tables 4 and 6 for a list of brain regions depicted in this figure.

**Table 4.** Categorical responses to object sounds specific to the blind. Object selective regions specific to the blind are depicted in Figure 4. For each region significant in the between-group contrasts (left-hand table), corresponding coordinates significant in the main effect in the blind are listed in the right-hand table. None of these regions were activated in the sighted, indicating that the between-group effects (blind > sighted) are driven by these regions being responsive only in the blind. Two regions (underlined in the left-hand table) showed selective deactivation in the sighted, thus contributing to the between-group effects observed in the R inferior OTC [34 −68 4] (z = 3.21) and in the L Middle Occipital G [−24 −84 6] (z = 3.25). Coordinates reported in this table are significant (p < 0.05 FWE) after correction over small spherical volumes (SVC) or over (*) the whole brain. K represents the number of voxels when displayed at p (uncorr)< 0.001. EB: early blind; SC: Sighted controls; V: voices; O: objects; SV: scrambled voices; SO: scrambled objects; L = left; R=Right; G=Gyrus; S=Sulcus, OTC = occipito-temporal cortex. Coordinates used for SVC are as follows (in MNI space): R Middle Occipital G: [44 −74 8] (Gougoux et al. 2009).

| Area | k | x<br>(mm) | y<br>(mm) | z<br>(mm) | Z | p | x<br>(mm) | y<br>(mm) | z<br>(mm) | Z | p |
| --- | --- | --- | --- | --- | --- | --- | --- | --- | --- | --- | --- |
| Between-group effects: [EB > SC] |  |  |  |  |  |  | Main effect in EB |  |  |  |  |
| **[O > SO]** |  |  |  |  |  |  | **[O > SO]** |  |  |  |  |
| L Inferior Occipital G | 7443 | -26 | -92 | 6 | 5.46 | 0.001* | -20 | -94 | 6 | 5.35 | 0.001* |
| L Middle/Inferior Occipital G |  | -36 | -82 | 0 | 5.45 | 0.001* | -32 | -84 | -4 | 5.14 | 0.003* |
| L Fusiform G |  | -36 | -68 | -14 | 4.86 | 0.011* | -34 | -68 | -18 | 5.82 | <0.001* |
| <u>R Inferior OTC</u> | 6075 | 38 | -66 | -4 | 4.75 | 0.018* | 38 | -66 | -6 | 4.54 | 0.039* |
| R Inferior Occipital G |  | 36 | -84 | 0 | 4.57 | 0.035* | 34 | -82 | -4 | 4.52 | 0.043* |
| R Fusiform S |  | 34 | -54 | -16 | 4.52 | 0.042* | 38 | -56 | -20 | 5.31 | 0.002* |
| **[O > V]** |  |  |  |  |  |  | **[O > V]** |  |  |  |  |
| L Middle Occipital G | 15494 | -26 | -92 | 4 | 5.57 | <0.001* | -24 | -94 | 8 | 5.39 | 0.001* |
| L Middle/Inferior Occipital G |  | -28 | -78 | -4 | 5.32 | 0.001* | -26 | -78 | -4 | 5.54 | 0.001* |
| L Superior Occipital G |  | -26 | -94 | 26 | 5.22 | 0.002* | -26 | -94 | 24 | 5.34 | 0.001* |
| **[O > SO] ∩ [O > V]** |  |  |  |  |  |  | **[O > SO] ∩ [O > V]** |  |  |  |  |
| L Middle Occipital G | 5582 | -26 | -92 | 6 | 5.46 | 0.001* | -24 | -94 | 6 | 5.31 | 0.001* |
| L Middle/Inferior Occipital G |  | -36 | -82 | -2 | 5.21 | 0.002* | -30 | -80 | -6 | 5.14 | 0.003* |
| L Lingual G |  | -20 | -74 | 0 | 4.49 | 0.047* | -20 | -72 | 2 | 4.83 | 0.013*** |
| R Inferior OTC | 4821 | 38 | -66 | -4 | 4.75 | 0.018* | 38 | -72 | 4 | 4.60 | 0.031* |
| R Inferior Occipital G |  | 36 | -80 | -2 | 4.40 | 0.001 | 36 | -76 | -4 | 4.15 | 0.002 |
| R Middle Occipital G |  | 40 | -86 | 10 | 4.29 | 0.001 | 40 | -82 | 10 | 3.75 | 0.008 |

On the lateral portion of the occipito-temporal cortex, these object selective responses specific to EB partially overlapped with shape-selective visual cortex localized in SC using the contrast [O>SO] (“Lateral occipital complex” or LOC, Malach et al. 1995) (Table 3), as also confirmed by a conjunction analysis (data not shown).

A conjunction (AND) analysis was conducted on the 2 sample t-tests [EB > SC] × [V > O] and [EB > SC] × [V > SV] in order to identify regions specifically activated in EB (relative to SC) for the processing of voices relative to both scrambled voices and objects. This analysis yielded no significant response, even at a very lenient threshold of p < 0.01 uncorrected. Considering each of these 2 sample-tests separately revealed that voices relative to scrambled voices [EB > SC] × [V > SV] elicited higher responses in EB in the fusiform gyrus bilaterally (Table 5). This effect was driven by the EB group (Table 5). In contrast, voices compared to objects [EB>SC] × [V>O] did not elicit any larger activation in EB relative to SC. The reverse group comparisons [SC > EB] did not reveal any region that was more strongly responsive in SC for voices relative to either objects or scrambled voices.

**Table 5.**
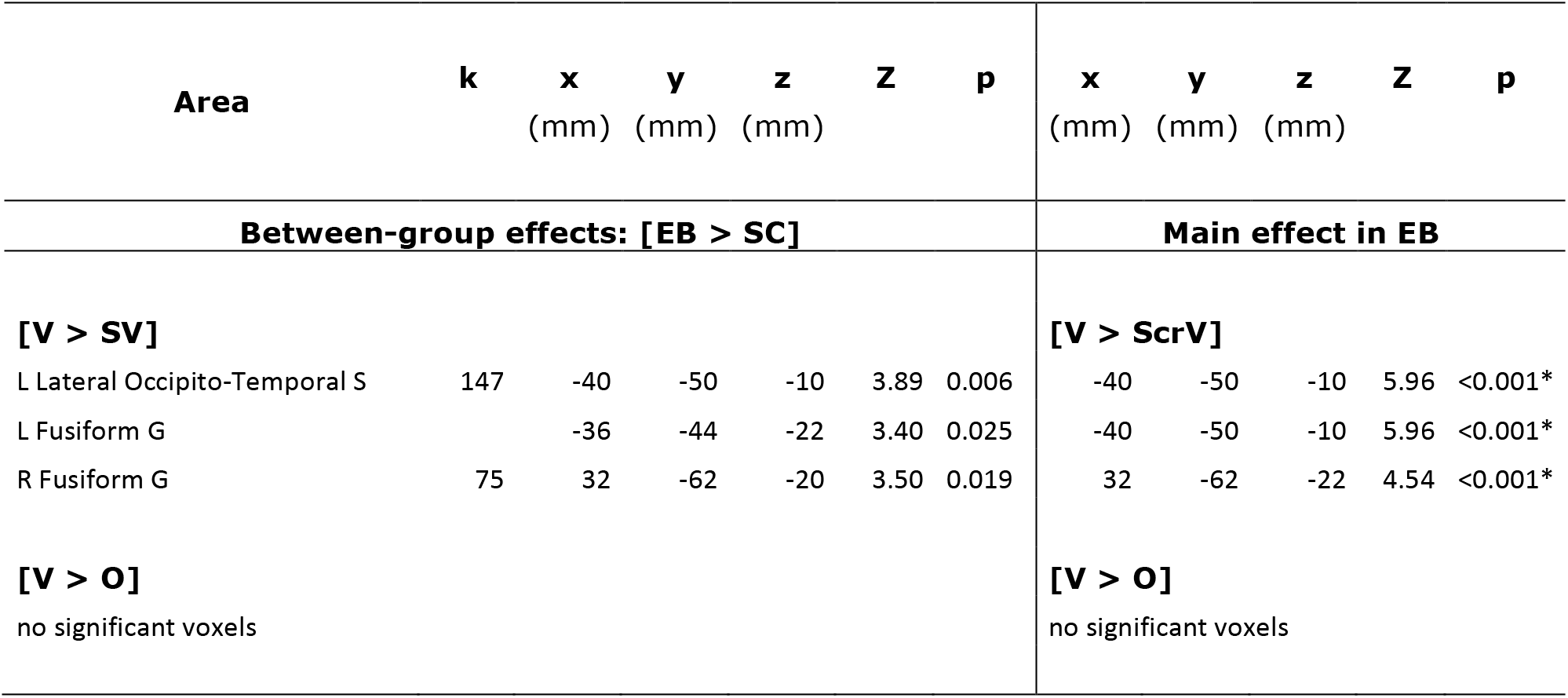
Voice-selective regions specific to the blind. These regions are depicted in Figure 5. For each region significant in the between-group contrast (left-hand table), corresponding coordinates significant in the main effect in the blind are listed in the right-hand table. None of these regions were activated or deactivated in the sighted, indicating that the between-group effects (blind > sighted) are driven by these regions being responsive only in the blind. Note that when contrasting voice to object sounds, no regions showed higher responses in the blind relative to the sighted. Coordinates reported in this table are significant (p < 0.05 FWE) after correction over small spherical volumes (SVC) or over (*) the whole brain. K represents the number of voxels when displayed at p (uncorr)< 0.001. EB: early blind; SC: sighted controls; V: voices; O: objects; SV: scrambled voices; SO: scrambled objects; L = left; R=Right; G=Gyrus; S=Sulcus. Coordinates used for SVC are as follows (in MNI space): L Fusiform/Inferior Temporal G: [−46 −48 −16] (Gougoux et al. 2009); R Fusiform G: [34 −52 −16] (Gougoux et al. 2009).

| Area | k | x<br>(mm) | y<br>(mm) | z<br>(mm) | Z | p | x<br>(mm) | y<br>(mm) | z<br>(mm) | Z | p |
| --- | --- | --- | --- | --- | --- | --- | --- | --- | --- | --- | --- |
| Between-group effects: [EB > SC] |  |  |  |  |  |  | Main effect in EB |  |  |  |  |
| **[V > SV]** |  |  |  |  |  |  | **[V > ScrV]** |  |  |  |  |
| L Lateral Occipito-Temporal S | 147 | -40 | -50 | -10 | 3.89 | 0.006 | -40 | -50 | -10 | 5.96 | <0.001* |
| L Fusiform G |  | -36 | -44 | -22 | 3.40 | 0.025 | -40 | -50 | -10 | 5.96 | <0.001* |
| R Fusiform G | 75 | 32 | -62 | -20 | 3.50 | 0.019 | 32 | -62 | -22 | 4.54 | <0.001* |
| **[V > O]**<br>no significant voxels |  |  |  |  |  |  | **[V > O]**<br>no significant voxels |  |  |  |  |

In other words, voice selective responses relative to both object sounds and scrambled voices were limited to the superior temporal sulci (auditory cortices) in both EB and SC with no evidence of crossmodal responses in the VOTC in either group (as also accounted by the individual data, data not shown).

### fMRI Results – Psychophysiological analyses

Psychophysiological interaction (PPI) analyses were computed to identify any brain regions showing a significant change in functional connectivity with specific seed areas as a function of experimental condition (O > V and V > O) and group (EB > SC) (Table 6).

**Table 6.**
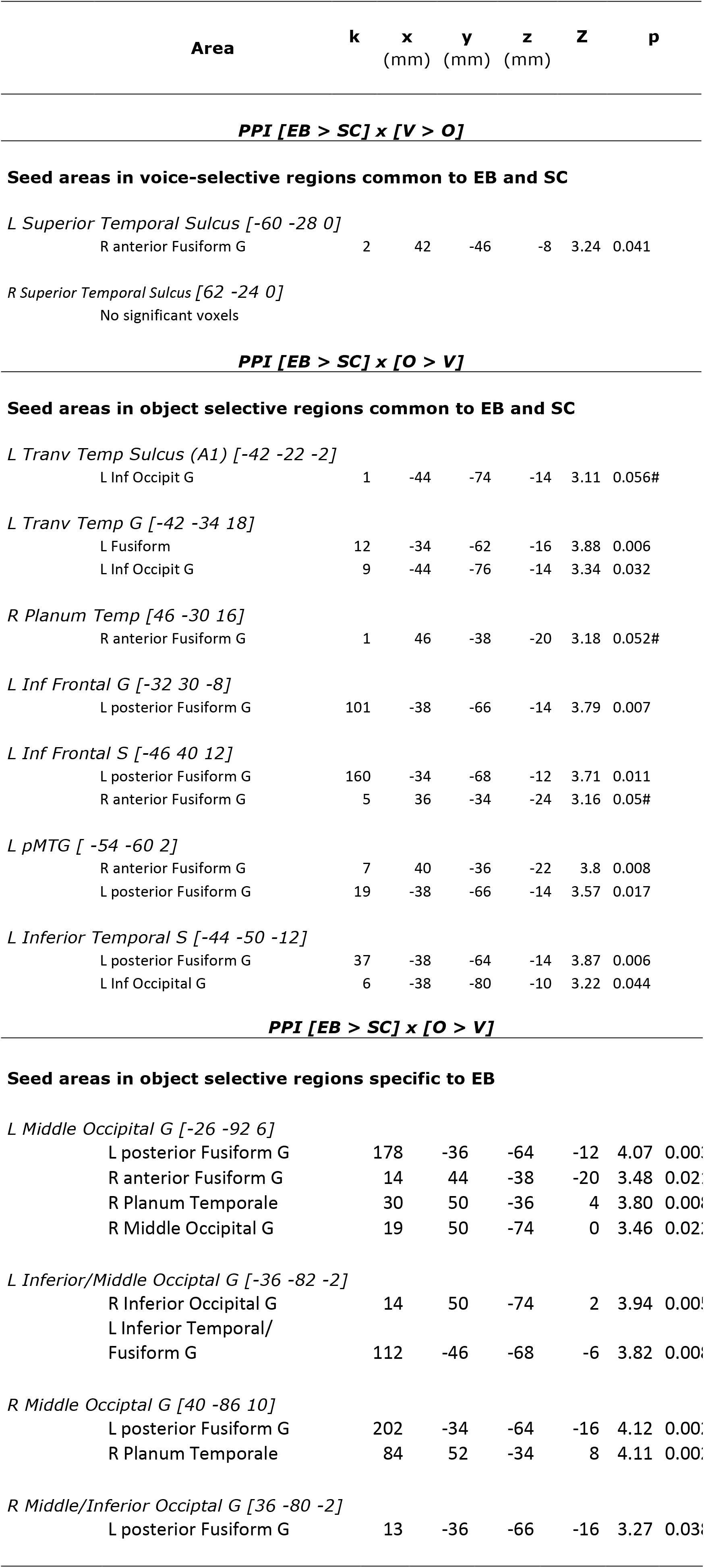
Regions showing increased functional connectivity with specific seed areas as a function of experimental condition (O > V and V > O) and group (blind > sighted). Seed areas are the ones resulting from the activation analyses (depicted in Figure 2A, 2B and 4A). **Regions showing increased connectivity with** these seed areas are listed in this table and are depicted in Figure 2D and Figure 4B. Coordinates reported in this table are significant (p < 0.05 FWE) after correction over small spherical volumes (SVC). Marginally significant clusters are indicated with (#). EB: early blind; SC: sighted controls; V: voices; O: objects; L = left; R=Right; G=Gyrus; S=Sulcus; pMTG: posterior middle temporal gyrus. Coordinates used for correction over small spherical volumes are as follows (in MNI space): R Fusiform G: [40 −36 −10] (Hölig et al. 2014); L Fusiform G: [−36 −63 −18] (Noppeney et al. 2003); L Inferior Occipital G: [−36 −81 −15] (Noppeney et al. 2003); R Planum Temporale: [52 −44 10] (Lewis et al. 2011); R Middle Occipital G: [44 −74 8] (Gougoux et al. 2009)

Among the two regions selectively responsive to voices in both groups (Figure 2A), the left superior temporal sulcus (−60 −28 0) displayed an increase in functional connectivity with the right fusiform gyrus in the blind during voice processing compared to object sounds processing (Figure 2D, a).

Among the regions that selectively responded to object sounds in both groups (Figure 2B), several seed areas located in auditory cortices showed a significant increase in functional connectivity with ventral occipito-temporal regions during object sounds relative to voice processing in blind relative to sighted subjects (Figure 2D, c-e). Notably, the left primary auditory cortex (−42 −22 −2) showed increased connectivity with the left inferior occipital gyrus (Figure 2D, c), the left transverse temporal gyrus (−44 −34 18) showed increased connectivity with the left inferior occipital gyrus and the left posterior fusiform gyrus (Figure 2D, d), while the right planum temporale (46 −30 16) showed increased connectivity with the right fusiform gyrus (Figure 2D, e). Regions located in the left inferior frontal gyrus (−32 −30 −8) and sulcus (−46 40 12) and those located in the left temporal cortex (left posterior middle temporal gyrus −54 −60 2, left inferior temporal sulcus −44 −50 −12), all showed an increase in functional connectivity with a circumscribed region located in the left posterior fusiform gyrus (Figure 2D, f-i). In addition, the left inferior frontal sulcus (−46 −40 −12) and the posterior middle temporal gyrus (−54 −60 2) showed an increase with the right anterior fusiform gyrus (Figure 2D, f-g), while the left inferior temporal sulcus (−44 −50 −12) showed an increase with the left inferior occipital gyrus (Figure 2D, h).

Among the reorganized occipital regions that showed a categorical response to object sounds only in EB (Figure 4A), all showed increased connectivity with a circumscribed region in the left posterior fusiform gyrus (Figure 4B, a-d). In addition, the left middle occipital gyrus (−26 − 92 −6) showed increased connectivity with the right anterior fusiform gyri, the right middle occipital gyrus and the right planum temporale (Figure 4B, a). The left inferior occipital gyrus (−36 −82 −2) showed increased functional connectivity with the right inferior occipital gyrus (Figure 4B, b), and the right middle occipital gyrus (40 −86 10) showed an increase in connectivity with the right planum temporale (Figure 4B, c).

## Discussion

The present study investigates how visual experience impacts on the neural basis of object sounds and voice processing. We used scrambled control sounds to control for low-level differences in the frequency spectrum between these categories of sounds and assess the contribution of low-level parameters to the categorical responses observed for objects sounds and voices.

### Double dissociation for object sounds and voices in the auditory temporal cortices

In both EB and SC, a double dissociation was identified in the auditory cortices between separate regions showing categorical responses to either object sounds or voices, suggesting that the cortical pathways for processing these 2 auditory categories are at least partially separable (Figure 2, Table 2). In line with previous work, categorical responses to sounds of objects were observed along the lateral fissure bilaterally (Giordano et al., 2013; Lewis et al., 2005; 2012; Lewis, Talkington, Puce, Engel, & Frum, 2011b) whereas categorical responses to voices were observed within bilateral superior temporal sulci (Belin et al., 2000; 2002; Belin, Fecteau, & Bédard, 2004). These findings are in line with the idea that the auditory system – like the visual one – hosts a domain-specific organization where distinct areas preferentially respond to different categories of complex environmental sounds such as voices, animal vocalizations, tools or musical instruments (Engel et al., 2009; Lewis et al., 2005; 2009; Patterson, Uppenkamp, Johnsrude, & Griffiths, 2002). These results are also in line with neuropsychological evidence demonstrating that lesions to portions of the temporal or temporoparietal cortex can lead to auditory agnosia, an impaired capacity to recognize complex natural sounds despite preserved speech comprehension and visual object recognition (for a review see Goll, Crutch, & Warren, 2010).

Our paradigm further allowed us to investigate the contribution of low-level parameters to these categorical responses. Our scrambling technique mainly preserved the frequency content of the sounds (Figure 1) while altering harmonic and phase-coupling content that is known to participate to the response in voice selective regions (Lewis et al., 2009). Here, higher responses in voice-selective areas were observed when contrasting scrambled voices and objects (SV > SO) (Table 2). This suggests that the spectral frequency content of voices strongly participates to the signal attributes that preferentially activate these regions. In other words, the selectivity observed for voices compared to objects in bilateral STS may emerge, at least partly, from the differential processing of low-level features that are typical of these two categories of sounds (Lewis et al., 2009; see Andrews, Clarke, Pell, & Hartley, 2010 for a similar interpretation in vision). In contrast, low-level parameters did not contribute to object categorical responses since no area in the brain was more largely responsive to scrambled objects than to scrambled voices.

Interestingly, no between-groups differences were observed for these domain-selective responses in temporal ‘auditory’ cortex, contrasting with previous findings (Gougoux et al., 2009; Lewis, Frum, Brefczynski-Lewis, Talkington, Walker, Rapuano, et al., 2011a). Based on a large sample of EB (n = 16) and controlling for the low-level properties of the auditory stimuli, the present findings therefore suggest that most of the brain reorganization ensuing early visual deprivation expresses outside of the temporal auditory cortices.

### Supramodal object representations in the left lateral and ventral occipito-temporal cortex

Beyond the auditory cortex, preferential responses to object sounds common to both EB and SC were found in left-lateralized inferior frontal and occipito-temporal regions including the posterior middle temporal, inferior temporal and fusiform gyrus (Fig 2B, Table 2). These left frontal and temporal regions have been previously associated with auditory object recognition (Lewis et al., 2004) and with semantic processing more generally, especially concerning concrete objects (Gold et al., 2006; Gough, Nobre, & Devlin, 2005; Sharp, Scott, & Wise, 2004; Wheatley, Weisberg, Beauchamp, & Martin, 2005, for a review see Martin, 2007). In fact, in the present study, these regions also responded selectively when sighted participants viewed pictures of objects (compared to both faces and scrambled objects) (Figure 3B, Table 3).

In previous studies, similar left fronto-temporal regions were found to be responsive in both early blind and sighted subjects in tasks involving action-related semantics (left inferior frontal [−51 30 3] and left posterior MTG [−63 −51 −6] Noppeney et al., 2003), sounds of tools (left pMTG [−51 −57 3] Lewis et al., 2005) heard names of tools (left pMTG [−50 −52 −3] Peelen et al., 2013) and places (left parahippocampal gyrus/fusiform [−28 −26 −21] C. He et al., 2013) as well as in sighted subjects viewing pictures of corresponding objects (He et al., 2013; Peelen et al., 2013).

The finding of preferential responses to “objects” independent of the input modality (visual and auditory) and of visual experience (since they are also observed in EB) in left occipito-temporal regions may suggest that these regions support a supra-modal organization of object representations (Bi et al., 2016; Fairhall & Caramazza, 2013). Because all sounds of objects in this study were highly recognizable, we speculate that these abstract representations were automatically activated when participants were listening to these familiar environmental sound-sources (see Lewis et al., 2004 for similar interpretation). In line with previous studies, we thus propose that the left occipito-temporal regions showing object selective responses in the present study contain an abstract representation of objects - such as the object’s meaning and semantics knowledge associated with it (Bi et al., 2016; Bracci, Cavina-Pratesi, Ietswaart, Caramazza, & Peelen, 2012; Fairhall & Caramazza, 2013; Kassuba et al., 2011; Lewis et al., 2004) - which develops independently of visual experience.

An alternative interpretation to supra-modality in left occipito-temporal regions is visual mental imagery: the latter could have driven responses to auditory stimuli in the sighted, as previously demonstrated for the tactile exploration of objects (Lacey, Flueckiger, Stilla, Lava, & Sathian, 2010). In other words, the possibility that similar activation patterns in EB and SC are related to different cognitive processes cannot be ruled out. For instance, the left occipitotemporal regions could be part of the visual sensory cortex in the sighted and be responsive in the auditory task because of visual imagery, while these same regions could have reorganized in the blind in order to support more abstract representations of objects and semantics. While no study to date can conclusively rule visual imagery out, we attempted to minimize this potential confound in our task by focusing the participant’s attention on the acoustical properties of the sounds.

Functional connectivity analyses performed on the left inferior frontal cortex and pMTG showed a unique connectivity pattern in EB, namely an increased task-related coupling with the left fusiform gyrus. These findings are remarkably similar to the ones reported by Noppeney et al. (2003). These authors found that the left inferior frontal cortex and left pMTG were activated in both early blind and sighted subjects during a semantic retrieval task on verbal material. Yet, functional connectivity analyses on both of these regions revealed increased coupling with left lateralized occipito-temporal areas only in the blind. Altogether, these findings suggest that early visual deprivation, although preserving the responsiveness of the inferior frontal cortex and left occipito-temporal cortex to object sounds (Figure 2B), impacts on these regions at the network level (Figure 2D). Worth noting, between-group differences in the connectivity profile of regions showing similar task-dependent activity level could support the idea that the cognitive processes underlying the recruitment of those regions partially differ between EB and SC.

### Crossmodal categorical responses to object sounds in posterior occipital cortex of the blind

A unique pattern of categorical responses to object sounds were found in EB within large portions of the occipital cortex, peaking in the middle and inferior occipital gyri bilaterally (Figure 4, Table 4). This suggests a posterior expansion of cortical function related to the representations of object’s sounds in the blind. These unique object selective responses in EB partially overlapped with portions of shape-selective visual cortex localized visually in the sighted (Malach et al., 1995) (Table 3). This runs counter to the idea that object-related responses in the occipital cortex of the blind rely solely on the processing of shape information conveyed by objects (either via touch or sensory substitution devices Amedi et al., 2007; 2010) since our stimuli and task did not involve shape processing. Interestingly, a trend for responses to object sounds in LOC was previously reported in 2 congenitally blind subjects when no imagery of shape was involved (Amedi et al., 2007). Together, these findings suggest that at least portions of LOC in early blind individuals contain representations of object sounds that are not related to shape and that these regions reorganize due to the lack of developmental vision since they do not activate in sighted individuals. Interestingly, crossmodal responses to object sounds in EB were mostly pronounced *outside* of visual regions with preferential responses to either shape (LOC, object pictures > scrambled objects) or object (objects > face) in the sighted: they extended more posteriorly in the occipital cortex (Figure 3 and Figure 4). Similar activation patterns with crossmodal responses extending posteriorly were reported in a previous study when congenitally blind subjects were involved in a tactile recognition task (Amedi et al., 2010), raising the intriguing possibility that cognitive processes underlying crossmodal occipital responses in the present study may not be specific to the auditory modality itself.

What may be the cognitive processes or representational format supporting the categorical responses to sounds of objects observed in EB in the present study? It has been proposed that environmental sounds that are perceived as “object-like”, such as those produced by automated machinery and man-made objects (as in the present study), share common acoustical features which may serve as low-level cues for their identification in a complex acoustic environment (Lewis et al., 2012). In the present study, none of the reorganized occipital regions showed stronger responses to scrambled objects compared to scrambled voices, running counter to the idea that categorical responses to object sounds are driven by low-level acoustic features that differentiate object sounds and voices (i.e. frequency spectrum). Rather, we argue that these occipital regions are an extension of more anterior occipito-temporal regions commonly responsive to sounds of objects in both SC and EB (Figure 3) and supporting abstract representations and semantics associated to the automatic processing of object’s meaning (see the discussion section on supramodality above). Several arguments account for this assumption. Object-selective crossmodal responses in EB were strongest in the left hemisphere, and in the vicinity of regions previously reported as being responsive when early blind subjects (compared to sighted subjects) process meaningful speech (sentences and word lists compared to nonsemantic sentences and non-word lists) (Bedny et al., 2011, Röder et al., 2002), generate semantically related verb to heard nouns (Amedi, Raz, Pianka, Malach, & Zohary, 2003; Burton, Diamond, & McDermott, 2003), and perform semantic decisions on heard nouns (Noppeney et al., 2003). Moreover, the functional connectivity pattern of these reorganized occipital regions in EB resembles the one observed for the left pMTG and inferior frontal cortex, that is, a systematic increased coupling with ventral occipito-temporal regions (inferior temporal/fusiform gyrus), mainly in the left hemisphere. Hence, we propose that the left pMTG showing object’s sound selectivity in both EB and SC on the one hand, and more posterior occipital regions showing preferential response to object’s sounds only in EB on the other hand, support similar functions, that is, an abstract representation of object semantics. While such representations are shared across modalities and populations in more anterior occipito-temporal regions, in line with a pure definition of “supramodal” representations, posterior occipital regions might support similar functions only in the early blind due to crossmodal plasticity.

Future studies directly comparing listening to object’s sounds versus listening to speech material would provide additional information to shed light on the mechanisms driving the occipital responses to object sounds observed in early blind subjects.

### No crossmodal categorical responses to voice in early blind or sighted subjects

In contrast to our observation of categorical responses to object’s sounds in the occipitotemporal cortex of the sighted and, to a larger extent, of the blind, no such categorical responses to voices were observed outside of the temporal auditory cortices in either group. This cannot be related to a lack of sensitivity of our paradigm to detect voice-selective responses because preferential responses to voices compared to both object sounds and scrambled voices were successfully identified in bilateral superior temporal sulci in both EB and SC (Figure 2A) and in each single participant (data not shown).

Rather, our findings suggest that different auditory functions are not equally susceptible to be supported by the occipital cortex in case of early visual deprivation. In the same vein, we have previously shown that the spatial processing of sounds preferentially activates right dorsal regions of the occipital cortex in early blind subjects whereas pitch processing of sounds does not (Collignon et al., 2011; Collignon, Lassonde, Lepore, Bastien, & Veraart, 2007). We conclude that preferential responses to voices over non-vocal auditory objects are confined to areas of the superior temporal sulci, even in case of early visual deprivation. Yet, early visual deprivation seems to affect these regions at the network level since functional connectivity analyses identified unique patterns of connectivity in EB between the left TVA and the right fusiform gyrus (Figure 2D (a)). This does not exclude the possibility that the VOTC supports identification of auditory objects in general – vocal and non-vocal – in the blind. For instance, in a recent fMRI study, Hölig et al. (2014) reported a voice (speaker) congruency effect in the right anterior fusiform gyrus of congenitally blind subjects, suggesting that this region may have reorganized to support person identification through the auditory modality in case of early visual deprivation (Hölig, Föcker, Best, Röder, & Büchel, 2014). Yet, the absence of another category of sounds prevents from concluding that this effect represents a categorical preference for voices, since a similar congruency effect could have been observed in the same region for other non-vocal sounds. In the present study, selective responses to voices over scrambled voices were found in bilateral fusiform gyri of the blind (Figure 5), about 3 cm more posteriorly than the region reported by Hölig et al. (2014). However, responses in these regions were also significantly larger for objects sounds compared to their scrambled counterpart and, if anything, significantly larger for objects sounds than voices.

**Figure 5.**
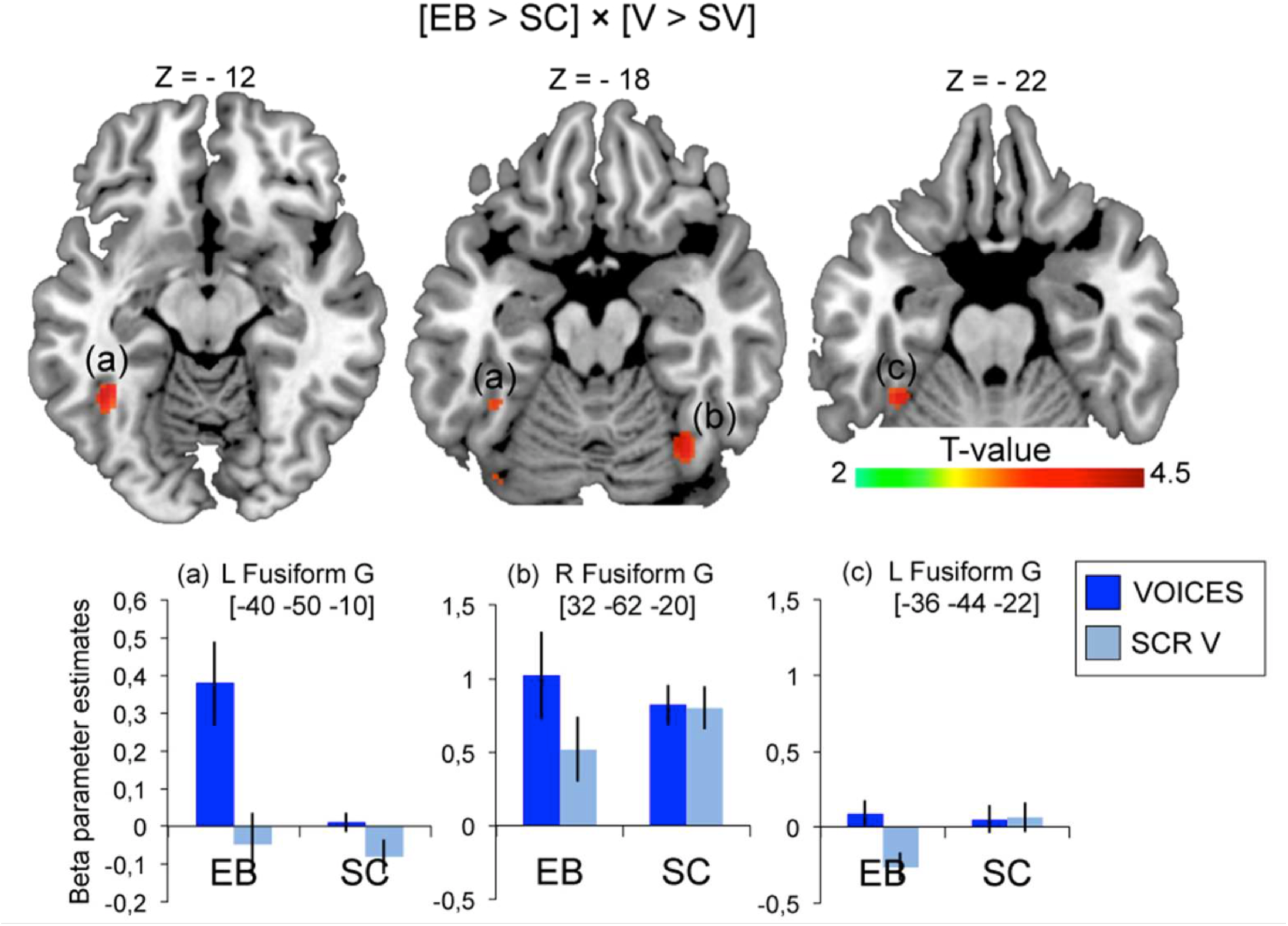
Regions showing stronger response to voices relative to scrambled voices specifically in the blind. For illustration purposes, activity maps are displayed at p(unc)< 0.001 with k > 10. Mean activity estimates (arbitrary units ± SEM) are plotted for voices and scrambled voices in significant peaks. EB = early blind; SC=sighted controls; L=left; R=right; S=sulcus; G=gyrus. See Table 5 for a list of brain regions depicted in this figure.

Similar conclusions of a lack of crossmodal reorganization of the face processing system in case of congenital blindness arise from a previous study investigating patterns of response elicited during tactile exploration of face masks and manmade objects in VOTC (Pietrini et al., 2004, see also Goyal, Hansen, & Blakemore, 2006). Category-related patterns of response in VOTC were found in sighted and blind for man-made objects (shoes and bottles) but not for face masks (Pietrini et al., 2004). Moreover, in the sighted, category-related patterns correlated across the visual and the tactile modality for manmade objects but not for faces. Based on these observations, the authors concluded that while objects’ representations might be supramodal in the VOTC, face representations are specific to vision. In the same vein, more recent studies reported overlapping responses to names of non-living objects in the VOTC of blind and sighted subjects (C. He et al., 2013; Peelen et al., 2013), while category-related responses to animals in the VOTC were only observed in the sighted and only with visually-presented material (C. He et al., 2013). It has thus been proposed that selectivity for non-living stimuli is multi-modal and independent of visual experience, while selectivity for living items, particularly in the lateral fusiform gyrus, is driven by visual stimulation only (Bi et al., 2016). Our findings of a lack of categorical responses to voices combined with preferential responses to objects’ sound in VOTC of the blind and the sighted are in agreement with this theoretical framework and suggest that regions supporting the representation of faces in the sighted brain do not transfer their preferential tuning to human voices in case of early blindness.

This lack of plasticity of the face recognition system is in line with the high degree of specialization (domain-specificity or modularity) of this system in typically developed individuals. Studies on the ontogeny of face recognition demonstrate impressive face recognition skills in newborns within a few days of birth (Johnson, Dziurawiec, Ellis, & Morton, 1991) and in monkeys raised without any exposure to faces (Sugita, 2008). Moreover, categorical neural responses to faces embedded among various non-face objects were recently identified in 4 months old babies (de Heering & Rossion, 2015). In the non-human primate brain, face responsive areas contain neurons responding selectively to faces (Desimone, 1991; Gross, Rocha-Miranda, & Bender, 1972; Tsao, Freiwald, Tootell, & Livingstone, 2006) and such areas have been demonstrated to be strongly interconnected and isolated from the rest of the visual recognition system at least (Moeller, Freiwald, & Tsao, 2008). Together, these characteristics of the face recognition system could come at the expense of generalization (to other domains) and plasticity. Some have proposed that the development of face recognition may be under high genetic control (Kanwisher, 2010). This assumption is supported by studies on families with hereditary prosopagnosia (Duchaine, Germine, & Nakayama, 2007; T. Grüter, Grüter, & Carbon, 2008; Schmalzl, Palermo, & Coltheart, 2008) and performance of monozigotic relative to dizigotic twins in a face memory task (Wilmer et al., 2010). In the same vein, Polk and collaborators (2007) found that genetics may play a larger role on neural activity patterns evoked by faces (and places) (Polk, Park, Smith, & Park, 2007) compared to the ones evoked by written peuso-words, the latter being more dependent on experience (J. Park, Park, & Polk, 2012; Polk et al., 2007; but see Pinel et al., 2014). Hence, different functional areas in the cortex may result from different neurodevelopmental mechanisms (Kanwisher, 2010). For example, while the visual word form area selectivity for word strings may emerge through pure learning-dependent mechanisms (Dehaene et al., 2010; S. He, Liu, Jiang, Chen, & Gong, 2009), face selectivity in the FFA may arise because “the specific instructions for constructing the critical circuits for face perception are in the genome” (Kanwisher, 2010). These different developmental mechanisms for defining functional areas might interact with sensory deprivation and therefore influence and constrain the process of crossmodal plasticity. In sum, the finding of crossmodal categorical responses to objects but not voices in the occipital cortex of early blind individuals suggests that crossmodal compensation in case of early visual deprivation depends on the neural systems investigated and on the neurodevelopmental mechanisms based on which these systems emerge.

## Acknowledgments

This work was supported by the Canada Research Chair Program (FL), the Canadian Institutes of Health Research (FL), the Belgian National Fund for Scientific Research (GD) and a European Research Council starting grant (MADVIS grant #337573) attributed to OC.

